# The Genetic History of France

**DOI:** 10.1101/712497

**Authors:** Aude Saint Pierre, Joanna Giemza, Matilde Karakachoff, Isabel Alves, Philippe Amouyel, Jean-François Dartigues, Christophe Tzourio, Martial Monteil, Pilar Galan, Serge Hercberg, Richard Redon, Emmanuelle Génin, Christian Dina

## Abstract

The study of the genetic structure of different countries within Europe has provided significant insights into their demographic history and their actual stratification. Although France occupies a particular location at the end of the European peninsula and at the crossroads of migration routes, few population genetic studies have been conducted so far with genome-wide data. In this study, we analyzed SNP-chip genetic data from 2 184 individuals born in France who were enrolled in two independent population cohorts. Using FineStructure, six different genetic clusters of individuals were found that were very consistent between the two cohorts. These clusters match extremely well the geography and overlap with historical and linguistic divisions of France. By modeling the relationship between genetics and geography using EEMS software, we were able to detect gene flow barriers that are similar in the two cohorts and corresponds to major French rivers or mountains. Estimations of effective population sizes using IBDNe program also revealed very similar patterns in both cohorts with a rapid increase of effective population sizes over the last 150 generations similar to what was observed in other European countries. A marked bottleneck is also consistently seen in the two datasets starting in the fourteenth century when the Black Death raged in Europe. In conclusion, by performing the first exhaustive study of the genetic structure of France, we fill a gap in the genetic studies in Europe that would be useful to medical geneticists but also historians and archeologists.

## INTRODUCTION

*Gallia est omnis divisa in partes tres* [*commentarii de bello gallico*^*1*^] was one of the earliest demographic description of antique France (known as Gaul). These three parts were Aquitania, in South West, with Garonne and the Pyrenees mountains as borders; Belgia in North West, following the Seine as Southern border; and finally what we know as Celtic Gaul, that spanned from the Atlantic Ocean to the Rhine River and Alps. A fourth part of the present-day French territory, already part of Romanized territories at this time, was Gallia Transalpina, a strip of lands from Italy to Iberia, with Alps and Cevennes mountains as northern border.

The area that was to be modern France was subject to successive population migrations: Western Hunter-Gatherers (15 kya), Neolithic farmers (7 kya) and later steppe Enolithic Age populations^2,3^, Celtic expansion, integration in Roman empire, Barbarian Great migrations, whose demographical importance remains to be assessed. France’s position in Europe, at the edge of the Eurasian peninsula, has made it not only the final goal of a large number of, potentially massive, migrations but also a place of transit either to the North (British Isles) or the South of Europe (Iberian Peninsula) and North Africa, as well as an important crossroad for trade and exchanges.

Before France became a single political entity, its territory was divided into various kingdoms and later provinces, which often displayed fierce independence spirit towards the central power. The pre-Roman Gaul was divided into politically independent territories. After the fall of Roman Empire, the modern French territory was divided into Barbarian Germanic kingdoms (Franks, Wisigoths and Burgunds). After a short period of reunification and extension into the Carolingian Empire (VII^th^ century), the weakening of the central power led to the reduction of the Occidental France at its western part and the rise of local warlords gaining high independence within the Kingdom itself. The feudality period created provinces that were close to independence, although nominally linked through the oath of allegiance to the King of France (Figure S1).

During centuries, in spite of important backlashes such as the Hundred Years War, the French Kings managed to slowly integrate the Eastern lands as well as Britanny, enforcing in parallel the central power until the French Revolution. However, every province kept displaying political, cultural and linguistic differences, which could have left imprints in the genetic structure of modern French populations.

Geographically, modern France is a continental country surrounded by natural borders: the Atlantic Ocean on the West side, the Channel Sea up North, mountains (Pyrenees and Alps) closing the South-West and East/South-East borders, as well as the Mediterranean Sea on the South side. The Eastern side has the Rhine as a natural border on less than 500 kilometers while the Northeastern borders shows no notable obstacle and exhibits a continuum with Germany and Belgium. This complex history is expected to have shaped the genetic make-up of the current French population and left some footprints in its genetic structure.

The study of the genetic structure of human populations is indeed of major interest in many different fields. It informs on the demographic history of populations and how they have formed and expanded in the past with some consequences on the distribution of traits. Genetic differences between populations can give insights on genetic variants likely to play a major role on different phenotypes, including disease phenotypes^4^. This explains the growing interest of geneticist for human population studies that aim at describing the genetic diversity and are now facilitated by the rich genetic information available over the entire genome. In the last decades, several studies were performed using genome-wide SNP data often collected for genome-wide association studies. These studies have first shown that there exist allele frequency differences at all geographic scales and that these differences increase with geographic distances. Indeed, the first studies have shown differences between individuals of different continental origins^5-7^ and then, as more data were collected and marker density increased, these differences were found within continents and especially within Europe^8,9^. Several studies have also been performed at the scale of a single country and have shown that differences also exist within country. This was for instance observed in Sweden, where Humphreys et al.^10^ reported strong differences between the far northern and the remaining counties, partly explained by remote Finnish or Norwegian ancestry. More recent studies have shown structure in the Netherlands^11^, Ireland^12^, UK^13^ or Iberian peninsula^14^. Previous studies of population stratification in France have examined only Western France (mainly *Pays de le Loi*re and Brittany) and detected a strong correlation between genetics and geography^15^. However, no study so far has investigated the fine-scale population structure of the entire France using unbiased samples from individuals with ancestries all over the country.

In this paper, we applied haplotype-based methods that have been shown to provide higher resolution than allele-based approaches^13^ to investigate the pattern of fine-scale population stratification in France. We used two independent cohorts, 3C and SUVIMAX with more than 2 000 individuals whose birthplace covered continental France and genotyped at the genome-wide level, to assess the genetic structure of the French population and draw inferences on the demographic history.

## MATERIAL AND METHODS

### Data on SU.VI.MAX & 3C studies

Genetic data were obtained from two French studies, SU.VI.MAX^16^ and the Three-Cities study^17^ (3C) with the idea to compare if, by analyzing them independently, concordant results would be obtained. Indeed, one major drawback of genetic inferences obtained on population samples is the fact that they can strongly depends on how individuals were sampled and which genetic markers were used. Here, by using data from two studies that sampled individuals using different criteria and genotyped them on different SNP-chip, we should be able to draw more robust inferences.

For every individual, information on places of birth was available, either the exact location (3C study) or the “*département*” (SU.VI.MAX). *Départements* are the smallest administrative subdivisions of France. There are a total of 101 French *départements* and 94 of them are located in continental France. These units were created in year 1 789, during French Revolution, partly based on historical counties.

#### 3C Study

The Three-City Study was designed to study the relationship between vascular diseases and dementia in 9 294 persons aged 65 years and over. For more details on the study, see http://www.three-city-study.com/the-three-city-study.php. Analyses were performed on individuals who were free of dementia or cognitive impairment by the time their blood sample was taken and who were previously genotyped^18^. The geographical locations of individuals were defined according to the latitude and longitude of their place of birth, declared at enrolment. Individuals with missing place of birth or born outside continental France were excluded. A total of 4 659 individuals were included in the present study.

#### SU.VI.MAX

The study was initiated in 1 994 with the aim of collecting information on food consumption and health status of French people. A subset of 2 276 individuals born in any of the 94 continental French *départements* was included in this study. The geographic coordinates of each *départements* were approximated based on the coordinates of the corresponding main city.

### Quality control

Quality control of the genotypes was performed using the software PLINK version 1.9^19,20^. 3C: raw genotype data were generated in the context of a previous study^18^ on Illumina Human610-Quad BeadChip. Following the recommendations from Anderson et al.^21^, individuals were removed if they had a call rate < 99%, heterozygosity level ± 3 standard deviations (SD) from the mean. Cryptic relatedness was assessed by estimating pi_hat (the IBD test implemented in PLINK^20^) in each dataset after doing LD-based pruning. Individuals related to another individual from the sample with an IBD proportion of 0.1875 or above were removed (only one individual was kept from each pair). As a final quality control to exclude outlier individuals from populations, we performed principle component analysis (PCA) using the smartpca software from the EIGENSOFT package version 6.0.1^22^ and removed outliers across the first 10 eigenvectors. The default procedure was used for outlier removals with up to 5 iterative PCA runs and, at each run, removing of individuals with any of the top 10 PCs departing from the mean by more than 6 standard deviations. SNPs in strong linkage disequilibrium (LD) were pruned out with PLINK 1.9 (described in PCA section). Outlier individuals were removed prior to performing further analyses. Applying all these QC filters led to the removal of 226 individuals. To avoid redundant information from individuals born at the same place, we randomly selected only one individual from each place of birth. A total of 770 individuals covering the 94 continental French “*départements*” were included. All samples failing sample-level QC were removed prior to performing SNPs QC. Markers were removed if they had a genotype-missing rate > 1%, a minor allele frequency < 1% or departed from Hardy–Weinberg proportion (P ≤ 10^-7^). After QC, there were 770 individuals and 490 217 autosomal SNPs.

#### SU.VI.MAX

Genotype data of the 2 834 samples were available from previous studies using different SNP chips: 1 978 with Illumina 300k/300kduo and 856 with Illumina 660W. Individuals with an unknown birthplace or a birthplace outside of continental France were removed, 1 416 samples were left. Two individuals were removed because of a call rate < 95%. IBD statistic, calculated in PLINK version 1.9, didn’t identify any related samples with a threshold of 0.1875. SNPs were removed if they had a genotype-missing rate > 2%, a minor allele frequency < 10 % or departed from Hardy–Weinberg proportion (P ≤ 10^-5^). After QC, there were 1 414 individuals and 271 886 autosomal SNPs.

### Population structure within France

#### Chromopainter/FineSTRUCTURE analysis

For investigating fine-scale population structure, we used Chromopainter version 2 and FineSTRUCTURE version 2.0.7^23^. Data were phased with SHAPEIT v2.r790^24^ using the 1000 genomes dataset as a reference panel. In the 3C dataset, we removed 932 of these SNPs because of strand issues prior to phasing. Files were then converted to Chromopainter format using the ‘impute2chromopainter2.pl’ script. Chromopainter outputs from the different chromosomes were combined with chromocombine to generate a final coancestry matrix of chunck counts for FineSTRUCTURE. For the FineSTRUCTURE run we sampled values after successive series of 10 000 iterations for 1 million MCMC iterations following 10 million “burn-in” iterations. Starting from the MCMC sample with the highest posterior probability among all samples, FineSTRUCTURE performed 100 000 additional hill-climbing moves to reach its final inferred state (See^13^ for details). The final tree was visualized in R with the help of FineSTRUCTURE and ‘dendextend’ libraries. We checked that the MCMC samples were independent of the algorithm’s initial position by visually comparing the results of two independent runs starting from different random seeds. Good correspondence in the pairwise coancestry matrices of the two runs indicates convergence of the MCMC samples to the posterior distribution. Without loss of generality, we used the first of these two runs in our main analysis.

#### Ancestry profiles of the French population & Spatial pattern of genetic structure EEMS

We used ADMIXTURE v1.3^25^ to estimate mixing coefficients of each individual. We performed runs for values of K between 2 and 10, with 5-fold cross-validation using the set of pruned SNPs, as described in the PCA analyses. To identify if cluster differences existed, we performed a one-way analysis of variance (ANOVA) on the admixture components, followed by *post hoc* pairwise comparisons.

We estimated an effective migration surface using the software EEMS^26^. We run EEMS with slightly different grids to investigate how/whether these changes affected the results. Plots were generated in R using the “rEEMSplots” package according to instructions in the manual. For both datasets the full set of SNPs was included. For more information on the specific pipeline, see Supplementary Data.

#### IBD-estimated population size

We estimated the recent effective population size with IBDNe^27^. IBDNe was run with the default parameters and a minimum IBD segment length of 4 cM (mincm=4). We used the default settings to filter IBD segments from IBDseq v. r1206 software package^28^. Breaks and short gaps in IBD segments were removed with the merge-ibd-segments utility program. For IBD detection, we varied the minimum IBD segment length in centiMorgan units by the mincm parameter (mincm argument) from the default value, 2 cM, to 8 cM. IBDNe analysis was applied on the whole SU.VI.MAX and 3C datasets as well as on the major subpopulations from fineSTRUCTURE clustering. Growth rates were calculated with the formula 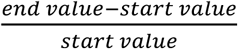. We assumed a generation time of 30 years, as assumed in the original paper.

#### Principal Component Analysis (PCA) and FST

Both PCA and FST analyses were carried out on a pruned set of SNPs in each dataset independently and using the smartpca tool in the EIGENSOFT program (v6.1.1)^22^. The pairwise FST matrix were estimated using the option ‘fsthiprecision=YES’ in smartpca. We calculated the mean FST between clusters inferred by FineSTRUCTURE as group labels. In each dataset, SNPs in strong LD were pruned out with PLINK in a two-step procedure. SNPs located in known regions of long range linkage disequilibrium (LD) in European populations were excluded from the analysis^29^. Then, SNPs in strong LD were pruned out using the ‘indep-pairwise’ command in PLINK. The command was run with a linkage disequilibrium r^2^=0.2, a window size of 50 SNPs and 5 SNPs to shift the window at each step. This led to a subset of 100 973 SNPs and 83 246 SNPs in the 3C and SU.VI.MAX datasets respectively. To evaluate the geographic relevance of PCs, we tested for the significance of association between the latitude and longitude of each *département* and PCs coordinates (‘cor.test’ function in R) using a Spearman’s rank correlation coefficient.

### Relation with neighbor European populations: 1000G and HGDP

We assembled SNP data matching either the SU.VI.MAX or the 3C genotype data (after quality control) with the European individuals from the 1000G phase 3 reference panel and from the Human Genome Diversity Panel data^30^ (HGDP, Illumina HuHap 650k), to generate four genome-wide SNP datasets analyzed independently.

The 1000G reference panel served as donor populations when estimating ancestry proportions. First, in order to define a set of donor groups from 1000G Europe (EUR), we used the subset of unrelated and outbred individuals generated in the study of Gazal et al.^31^. Four European populations were considered: British heritage (GBR, n=85 and CEU, n=94), Spain (IBS, n=107) and Italy (TSI, n=104). These 390 Europeans individuals were then combined with individuals from both datasets independently resulting in a set of 484 874 common SNPs with 3C and a set of 232 148 common SNPs with SU.VI.MAX. The filtered datasets (after pruning) included 1 160 individuals genotyped on 100 851 SNPs in the 3C Study and 1 804 individuals genotyped on 64 653 SNPs in SU.VI.MAX. We inferred European ancestry contributions in France using the novel haplotype-based estimation of ancestry implemented in SOURCEFIND^32^. SOURCEFIND has been shown to give a greater accuracy than the usual Non-Negative least squares regression for inferring proportion of admixture but because it is recommended to use homogeneous donor groups, we ran FineSTRUCTURE on the four European populations defined above and selected the level of clustering describing the main features of the donor populations. These European donor groups served as reference in SOURCEFIND. We performed analysis of variance (ANOVA) on French admixture component per cluster group to identify whether cluster differences existed.

Additional analyses combining the European participants of the HGDP panel were carried out in order to estimate the contribution of Basque population of our South West clusters. A total of 160 European HGDP participants were included from 8 populations: Adygei (n=17), French-Basque (n=24), French (n=29), Italian (n=13), Italian from Tuscany (n=8), Sardinian (n=28), Orcadian (n= 16) and Russian (n=25). Using the same procedure for merging panels, the filtered datasets (after pruning) included 930 individuals and 93 938 SNPs in the 3C Study and 1 574 individuals and 57 775 SNPs in SU.VI.MAX.

## RESULTS

### Chromopainter/FineSTRUCTURE analysis reveals consistent fine-scale genetic stratification within France

Results of FineSTRUCTURE analysis reveal fine-scale population patterns within France at a very fine level that are very consistent in the two datasets (Figure 1). FineSTRUCTURE identified respectively 17 and 27 clusters in 3C and SU.VI.M.AX, demonstrating local population structure (Figure S2). Even though the sampling distributions of individuals varies slightly between datasets both analyses show very concordant partitions with a broad correlation between clusters and geographic coordinates. The major axis of genetic differentiation runs from the south to the north of France.

**Figure 1:**
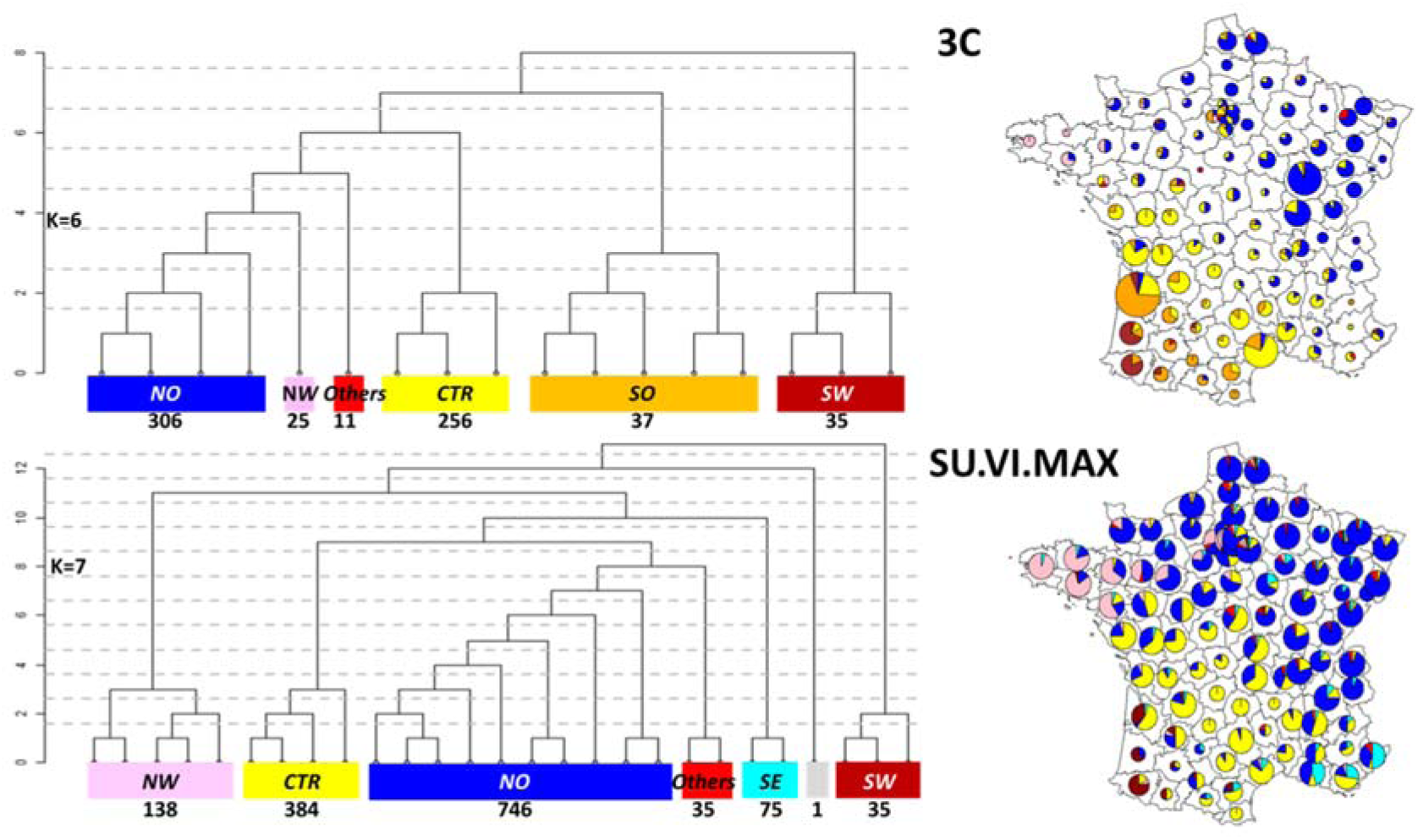
FineSTRUCTURE clustering of the 3C Study (770 individuals) and SU.VI.MAX (1 414 individuals) pooled in 6 and 7 clusters respectively. Left side shows the tree structure and right side shows by *département* pie charts indicating to which of the six clusters the individuals belong to.

In both datasets, the coarsest level of genetic differentiation (i.e. the assignment into two clusters) separates the south-western regions from the rest of France (Figure S3 and S4). Next levels of tree structures slightly differ between the two datasets but converge into a common geographic partitions at *k*=6 clusters in 3C and *k*=7 in SU.VI.MAX (Figure 2). The clusters are geographically stratified and were assigned labels to reflect geographic origin: the South-West SW for the dark-red cluster, the South (SO) for the orange cluster, the Centre (CTR) for the yellow cluster, the North-West (NW) for the pink cluster, the North (NO) for the blue cluster and the South-East (SE) for the cyan cluster. In each dataset, one cluster (labelled “*Others*” and coloured in red) included individuals geographically dispersed over France. Furthermore, one cluster identified in SU.VI.MAX included only one individual and was removed in further analysis so that *k*=7 also resumed to 6 clusters in SU.VI.MAX. At this tree level of 6 clusters, individuals from the NO, NW and CTR clusters are clearly separated in the two datasets. The SW cluster and part of the SO cluster in 3C match geographically the SW cluster identified in SU.VI.MAX while the SE subgroup was not detected in the 3C. This might be explained by differences in the geographic coverage between the two studies especially in the south of France. Indeed, SU.VI.MAX has a better coverage of the south-east whereas 3C lacks data from this region and the reverse is true for the south-west. In the two datasets, two large clusters (CTR and NO) are found that cover most of the central and northern France. Notably, even at the finest level of differentiation (17 and 27 clusters in 3C and SU.VI.MAX respectively), these clusters remain largely intact.

**Figure 2:**
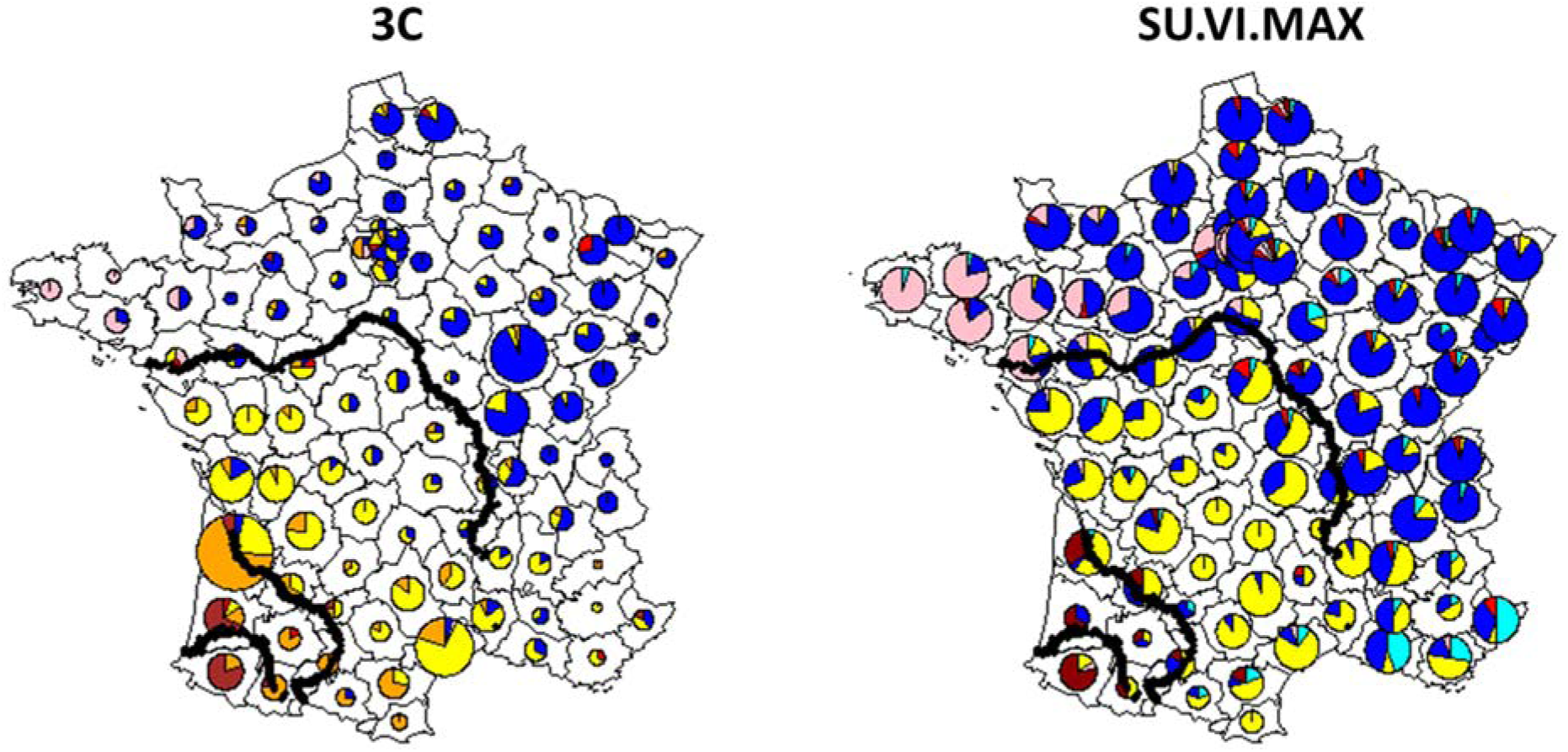
Pie charts indicating the proportion of individuals from the different “départements” assigned to each cluster. Results are reported for the partition in 6 clusters obtained by running FineSTRUCTURE in the 3C dataset (left) and in SU.VI.MAX (right) independently. Geographic coordinates of three rivers of France are drawn in black: Loire, Garonne and Adour from north to south.

The broad-scale genetic structure of France in six clusters strikingly aligns with two major rivers of France, “La Garonne” and “La Loire” (Figure 2). At a finer-scale, the “Adour” river partition the SW to the SO cluster in the 3C dataset. The mean FST between clusters inferred by FineSTRUCTURE (Table S1 and S2) are small, confirming subtle differentiation. In both datasets, the strongest differentiation is between the SW cluster and all other regions. These FST values vary from 0.0016 with the SO cluster to 0.004 with the NW cluster in the 3C dataset and from 0.0009 with the CTR cluster to 0.0019 with the NW cluster in SU.VI.MAX. Finally, besides this subtle division, genetic differentiation within France is also due to isolation by distance as shown by the gradient exhibited on the values of the 1^st^ component of the PCA (Figure S5).

### Different genetic ancestry profiles that could have been shaped by gene flow barriers

Results obtained by using ADMIXTURE corroborate the FineSTRUCTURE analysis with the SW cluster been the most different from the other groups (Figure S6 and S7). At k=2, the SW cluster shows a light blue component that is significantly less frequent in the other groups (ANOVA *post-hoc* tests, p-value<10^-6^) (Figure S8). In the 3C dataset, the proportion of light blue tends to decrease gradually from the south-western (SW) part of France to the centre of France (CTR) to finally remain similar in the north of France (NO, NW and *Others*). In SU.VI.MAX, the proportion of light blue component tends to discriminate the north from the south of France (Figure S8). For k=3, a third major component can be defined, the light green ancestry. In the 3C Study this component is predominant in the north of France (NW and NO clusters) and almost absent in the SW while in SU.VI.MAX this component is predominant in the SE and minimal in the extreme west of France (NW and SW). At k=6, both datasets highlight the differentiation of the SW and the NW cluster from the others clusters.

We performed EEMS analysis in order to identify gene flow barriers within France; i.e; areas of low migrations. We varied the number of demes from 150 to 300 demes and selected a grid of 250 demes showing good concordance between datasets (Figure S9). In both datasets, we identified a genetic barrier around the south-west region (Figure 3). This barrier mirrors the first division in the FineSTRUCTURE. The plots also reveal a gene flow barrier around Bretagne in the North-West and along the Loire River, which covers the separation of the North cluster. Finally, another barrier is also present on the South-East side that roughly corresponds to the location of the Alp Mountains at the border with Northern Italy.

**Figure 3:**
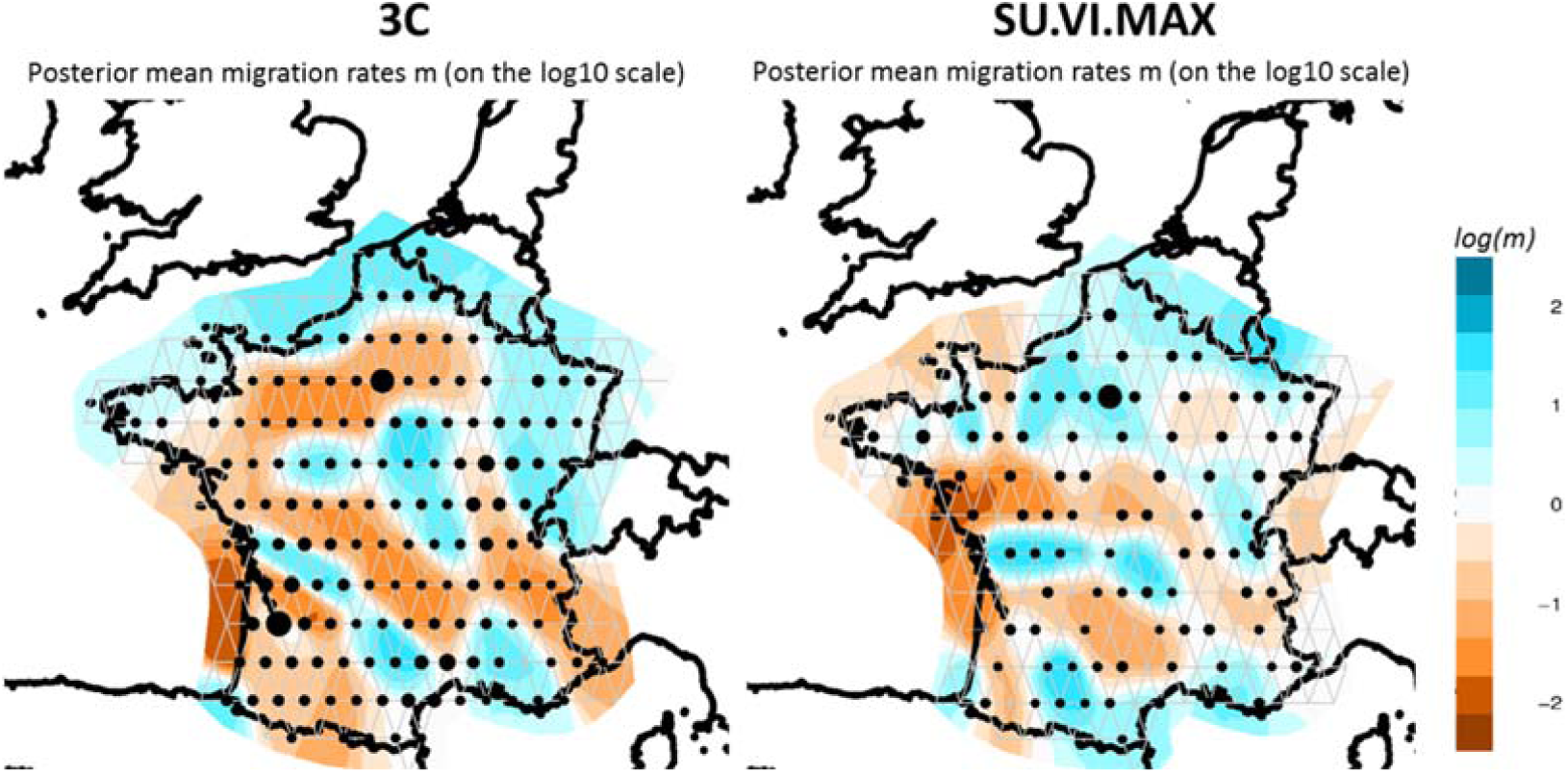
Estimated effective migration surfaces of France obtained from EEMS on the 3C (left) and SU.VI.MAX (right) datasets. The colour scale reveals low (blue) to high (orange) genetic barriers between populations localized on a grid of 250 demes. Each dot is proportional to the number of populations included.

### IBD-derived demographic inferences reveal a rapid expansion over the last 150 generations

Demographic inferences based on IBD patterns in the two datasets were also very concordant. We observed a very rapid increase of the effective population size, Ne, in the last 150 generations (Figure S10). The growth rate was 121% (0.8% per generation) in SUVIMAX and 100% (0.7% per generation) in 3C. This is in accordance with previous observations^33^ which reports a very rapid increase of human population size in Europe in the 150 last generations. However, the increase of Ne was not constant over time and a decrease of Ne is observed in both datasets in-between generations 12-22 (from ∼1 300 to 1 700). The growth rates in the period preceding and the period following this decrease were rather different. These growth rates were respectively of 3.1% and 2.4% per generation in 3C and SU.VI.MAX in the last 12 generations and of 0.4% and 0.6% per generation in the first period (22 to 150 generations ago). In-between these two periods, a bottleneck could be detected that could reflect the devastating Black Death. This decrease in Ne seems to affect mainly the Northern part of France (Figure S11). However, this result was not robust to change in parameters: the bottleneck effect was no longer seen when longer IBD chunks were used for Ne estimation (Figure S12).

### Different contributions of British and Basque heritage in the six French genetic clusters

To study the relationship between the genetic clusters observed in France and neighbor European populations, we combined our two datasets with the 1000G European dataset. As a first step, we run FineSTRUCTURE on the 1000G European populations excluding Finland and found that they could be divided into 3 donor groups as CEU and GBR clustered together (British heritage) (Figure S13). We estimated European ancestry contributions in France with SOURCEFIND and reported the total levels of ancestry proportions for each individual grouped by cluster (Figure 4). We observed similar patterns of admixture between datasets. The proportion of each admixture component from neighboring European countries was significantly different between the six FineSTRUCTURE clusters in both the 3C and SU.VI.MAX datasets (ANOVA, p-value<10^-16^). As expected, the British heritage was more marked in the north than in the south of France where, instead, the contribution from southern Europe was stronger. The overall contribution from British heritage was substantially higher in the NW than in the NO cluster (76% vs 64% in the 3C and 72% vs 63% in SU.VI.MAX). TSI was contributing to the SE cluster while IBS was mainly contributing to the SW cluster, which again was very coherent with the geographic places of birth of individuals. In both dataset, SW had the highest proportions of IBS component. Part of this IBS component could in fact reflect a Basque origin as shown on the PCA plot obtained when combining 3C, SU.VI.MAX and HGDP European dataset (Figure S14). This trend is even more pronounced in the 3C where few individuals are grouped together with Basque individuals in the first three dimensions. This SW region also corresponds to the “Aquitaine” region described by Julius Caesar in his “Commentari de Bello Gallico”^1^ (Figure S1).

**Figure 4:**
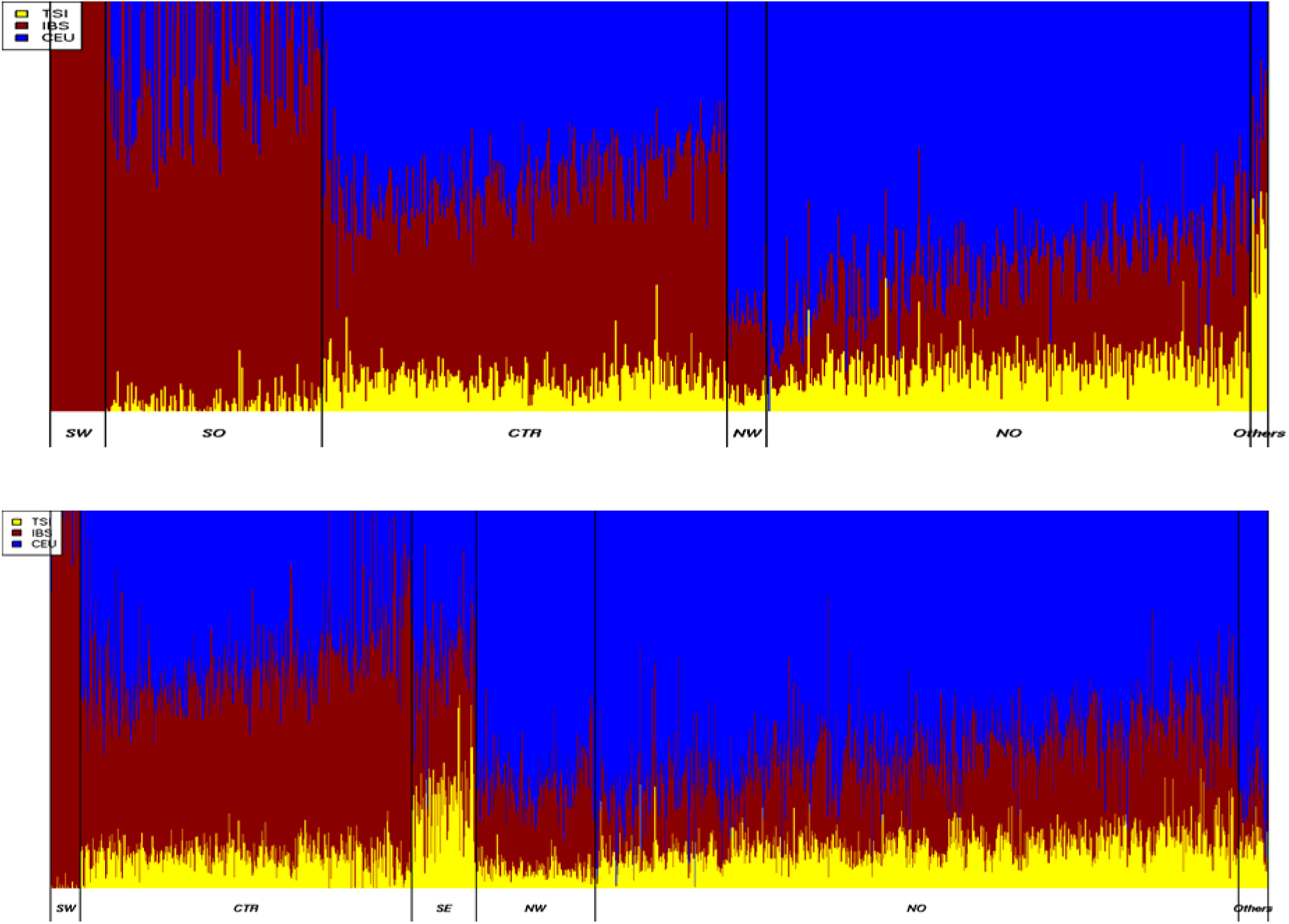
Ancestry profiles from the three neighbouring European populations inferred by SOURCEFIND in the French 3C (top) and SU.VI.MAX individuals (bottom) datasets. In each cluster, individuals are ordered according to the latitude of their reported birth place.

## DISCUSSION

In this paper, we have studied the genetic structure of France using data from two independent cohorts of individuals born in different regions of France and whose places of birth could be geolocalized. Modern France has a strategic location at the western-most part of Europe and on migration ways from and towards South and North of Europe. Studying its genetic structure is thus of major interest to gain insight on the peopling of Europe. To date however, no exhaustive study had been conducted on the French genetic make-up and our work was intended to fill this gap.

The French genomes were found to map at their expected position in between Nordic (British and CEU), Italian and Spanish genomes from 1000 genomes project. Within France, correlations were detected between genetic data and geographical information on the individual’s place of birth. Although the relation seems linear – reflecting isolation by distance process – we also observed that, based on genotype patterns, the population can be divided into subgroups, which match geographical regions and are very consistent across the two datasets.

An important division separates Northern from Southern France. It may coincide with the von Wartburg line, which divides France into “*Langue d’Oïl*” part (influenced by Germanic speaking) and “*Langue d’Oc*” part (closer to Roman speaking) – Figure S15. This border has changed through centuries and our North-South limit is close to the limit as it was estimated in the IXth century^34,35^. This border also follows the Loire River, which has long been a political and cultural border between kingdoms/counties in the North and in the South (Figure 3).

Regions with strong cultural particularities tend to separate. This is for example the case for Aquitaine in the South-West which duchy has long represented a civilization on its own. The Brittany region is also detected as a separate entity in both datasets. This could be explained both by its position at the end of the continent where it forms a peninsula and, by its history since Brittany has been an independent political entity (Kingdom and, later, duchy of *Bretagne*), with stable borders, for a long time^36^.

The extreme South-West regions show the highest differentiation to neighbor clusters. This is particularly strong in 3C dataset, where we even observe an additional cluster. This cluster is likely due to a higher proportion of possibly Basque individuals in 3C, which overlap with HGDP Basque defined individuals. The FST between the south-west and the other French clusters were markedly higher than the FST between remaining French clusters. In 3C these values are comparable to what we observed between the Italian and the British heritage clusters (FST=0.0035). Similar trends are observed in SU.VI.MAX even though the level of differentiation with the SW was weaker.

We also observe that the broad-scale genetic structure of France strikingly aligns with two major rivers of France “La Garonne” and “La Loire” (Figure 3). At a finer-scale, the “Adour” river partition the SW to the SO cluster in the 3C dataset.

While historical, cultural and political borders seem to have shaped the genetic structure of modern-days France, exhibiting visible clusters, the population is quite homogeneous with low FST values between-clusters ranging from 2.10^-4^ up to 3.10^-3^. We find that each cluster is genetically close to the closest neighbor European country, which is in line with a continuous gene flow at the European level. However, we observe that Brittany is substantially closer to British Isles population than North of France, in spite of both being equally geographically close. Migration of Britons in what was at the time Armorica (and is now Brittany) may explain this closeness. These migrations may have been quite constant during centuries although a two waves model is generally assumed. A first wave would have occurred in the X^th^ century when soldiers from British Isles were sent to Armorica whereas the second wave consisted of Britons escaping the Anglo-Saxon invasions^37^. Additional analyses, on larger datasets may be required to discriminate between these various models.

Studying the evolution of French population size based on genetic data, we observe a very rapid increase in the last generations. This observation is in line with what has been seen in European populations^33^. We also observe, in most cases, a depression during a period spanning from 12 to 22 generations ago. This may correspond to a period spanning from 1 300 to 1 700. Indeed, this period was characterized by a deep depression in population size due to a long series of plague events. While the population size in kingdom of France was estimated to be 20 Million in 1 348, it dropped down to 12 Million in 1 400, followed by an uneven trajectory to recover the 20 Million at the end of Louis XIV^th^ reign (1 715)^38^.

However, the decrease we observe in the genetic data does seem to affect mainly the Northern part of France, and for instance is mainly observed in the NO cluster. We see no reason for this trend based on historical records (Figure S16) except perhaps the last plague epidemics in 1 666 - 1 670 that was limited to the North of France. Alternatively, a more spread population in the South (which is in general hilly or mountainous) may explain a lower impact of these dramatic episodes. Plague is expected to have had a very strong impact on the population demography in the past as some epidemics led to substantial reduction in the population sizes^39^. However, we could not detect in our data any footprint of the Justinian plague (541-767 PC) although, according to historical records it had a major impact on the population at that time. This may be due to difficulty to estimate population changes in ancient times, deeper than 50-100 generations, especially in presence of more recent bottleneck and given our reduced sample sizes in some of the groups and IBD resolution power. We expect that increasing sample size especially for the FineSTRUCTURE subgroups with small sample sizes will help getting more detailed information farther in the past.

Identification of genetic structure is important to guide future studies of association both for common, but more importantly, for rare variants^40^. In the near future, interrogating the demographical history of France from genetic data will bring more precise results thanks to whole genome sequencing that, along with new methods, could allow testing formal models of demographic inference.

## ACKNOWLEDGMENTS

Part of this work was supported by the French National Research Agency (FROGH: ANR-16-CE12-0033) and the European Union via the Marie Skłodowska-Curie actions (PRESTIGE-2017-4-0018).

