## Supplemental Data for "The Genetic History of France"

### SUPPLEMENTARY DATA

#### Effective migration surface EEMS

We draw polygons around France with the online Google Maps tool (http://www.birdtheme.org/useful/v3tool.html). Depending on their location, several populations can be included in one deme, whose size increases accordingly. We ran the Markov chain Monte Carlo 5 times with different random seeds, each time with 9.9 million burn-in and 10 million regular iterations, thinning every two hundred iterations. For each deme, we chose the chain with the highest final log-likelihood, and started a second round of EEMS chains using this chain as a starting point and 1,000,000 additional sampling iterations thinning every 9,999 iterations. The dissimilarity between observed versus fitted deme pairs show a general trend with some deviation and the log-posterior trace of the replicate MCMC chains (Figure S9) show convergence of the independent EEMS runs.

### SUPPLEMENTARY TABLES

| $F_{ST}$ in 3C | SO | CTR | NW | NO | IBS | TSI | CEU+GBR |
|---|---|---|---|---|---|---|---|
| SW | 0.0016 | 0.0029 | 0.0040 | 0.0035 | 0.0022 | 0.0049 | 0.0047 |
| SO | | 0.0002 | 0.0013 | 0.0006 | 0.0003 | 0.0019 | 0.0016 |
| CTR | | | 0.0009 | 0.0002 | 0.0006 | 0.0016 | 0.0012 |
| NW | | | | 0.0006 | 0.0019 | 0.0033 | 0.0005 |
| NO | | | | | 0.0010 | 0.0017 | 0.0006 |
| IBS | | | | | | 0.0014 | 0.0023 |
| TSI | | | | | | | 0.0035 |

Table S1: $F_{ST}$ table between the French 3C clusters and the 1000G European clusters inferred by FineSTRUCTURE. Mean $F_{ST}$ statistics are estimated using EIGENSOFT.

| $F_{ST}$ in SU.VI.MAX | CTR | SE | NW | NO | IBS | TSI | CEU+GBR |
|---|---|---|---|---|---|---|---|
| SW | 0.0009 | 0.0015 | 0.0019 | 0.0014 | 0.0007 | 0.0028 | 0.0026 |
| CTR | | 0.0004 | 0.0007 | 0.0002 | 0.0005 | 0.0016 | 0.0012 |
| SE | | | 0.0013 | 0.0004 | 0.0006 | 0.0007 | 0.0017 |
| NW | | | | 0.0004 | 0.0017 | 0.0030 | 0.0005 |
| NO | | | | | 0.0010 | 0.0017 | 0.0006 |
| IBS | | | | | | 0.0015 | 0.0023 |
| TSI | | | | | | | 0.0035 |

Table S2: $F_{ST}$ table between the French SU.VI.MAX clusters and the 1000G European clusters inferred by FineSTRUCTURE. Mean $F_{ST}$ statistics are estimated using EIGENSOFT.

**SUPPLEMENTARY FIGURES**

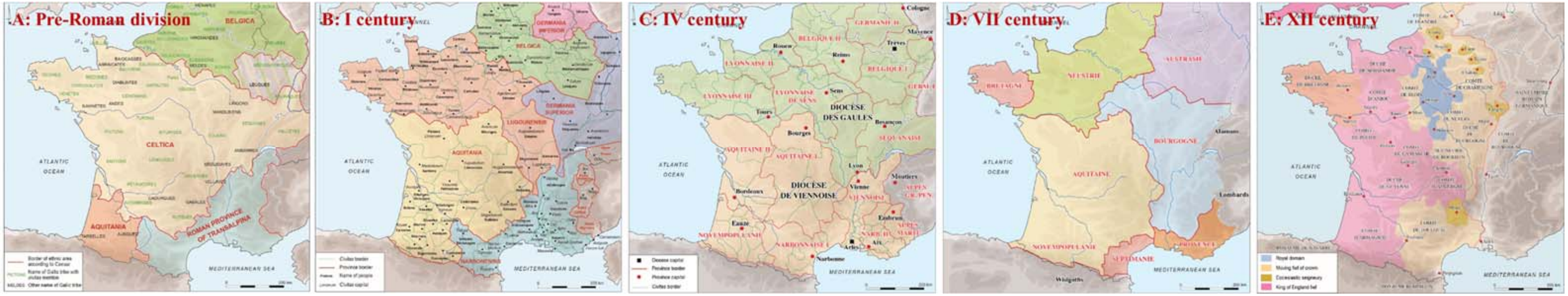


Figure S1: a/ Map of the territorial divisions of Gaul based on the description that Caesar made at the time of his conquest, between 58 and 51 BC (according to Fichtl. St., Les peuples gaulois. IIIe-Ier siècles av. J.-C., Paris, Errance, 2004, p. 10 et 54). b/ Map of the provinces and civitates of roman Gallia and Germania at the end of 1th c. AD. c/ Map of the provinces and civitates of roman Gallia and Germania at the end of 4th c. AD. d/ Map of political territories in the 7th c. E: Map of France in the 12th c. (according to Burnouf J., Archéologie médiévale en France. Le second Moyen Âge (XIIe-XVIe siècle), Paris, La Découverte, 2008, p. 98). Historical maps have been plotted by © M. Monteil, Nantes University, UMR 6566 CReAAH.

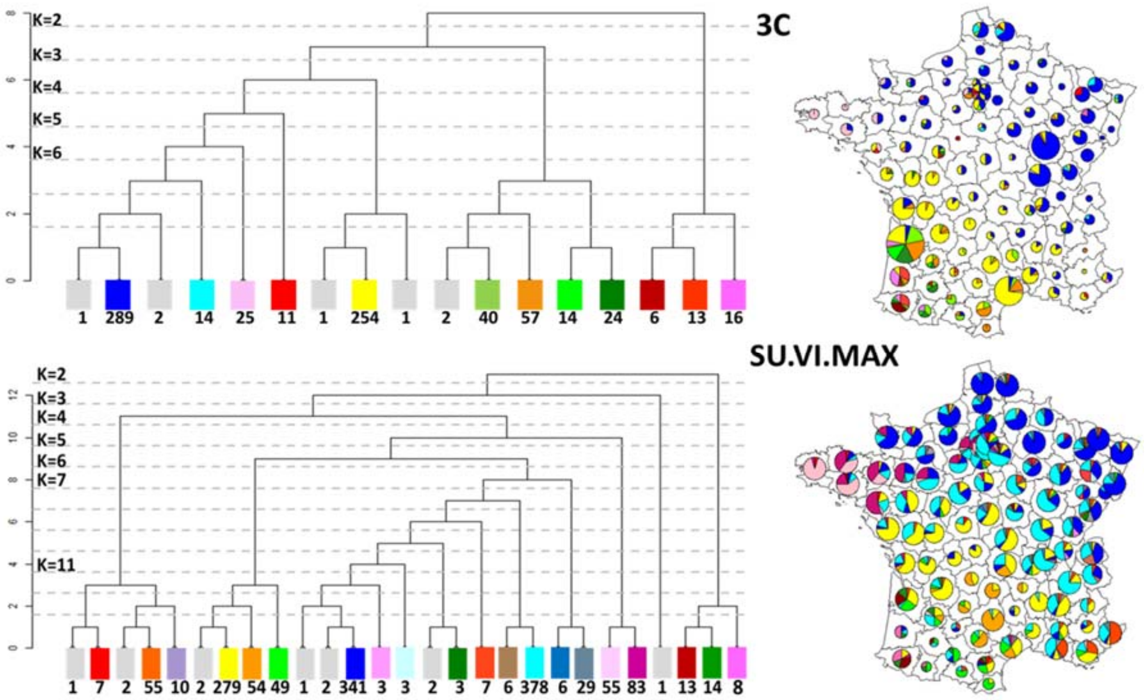


Figure S2: FineSTRUCTURE clustering of the 3C individuals (770 individuals, top) pooled in 17 clusters and SU.VI.MAX individuals (1,414 individuals, bottom) pooled in 27 clusters. Left side: tree structure. Right side: birth location of individuals coloured according to their assigned cluster. Clusters with less than 3 individuals are coloured in grey.

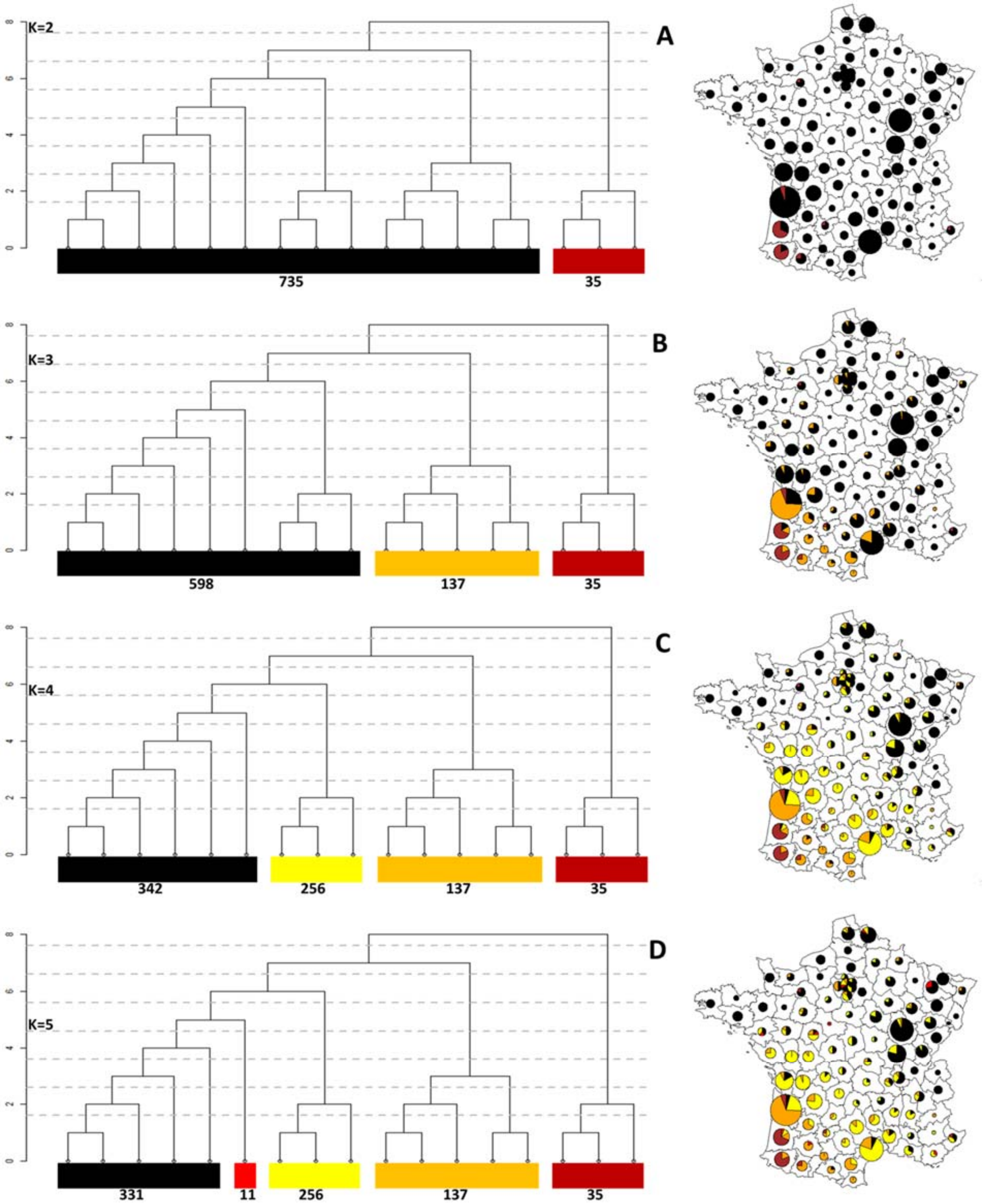


Figure S3: Genetic clusters in the 3C data inferred by the FineSTRUCTURE analysis at levels of the hierarchical clustering varying from k=2 (A), k=3 (B), k=4 (C) and k=5 (D).

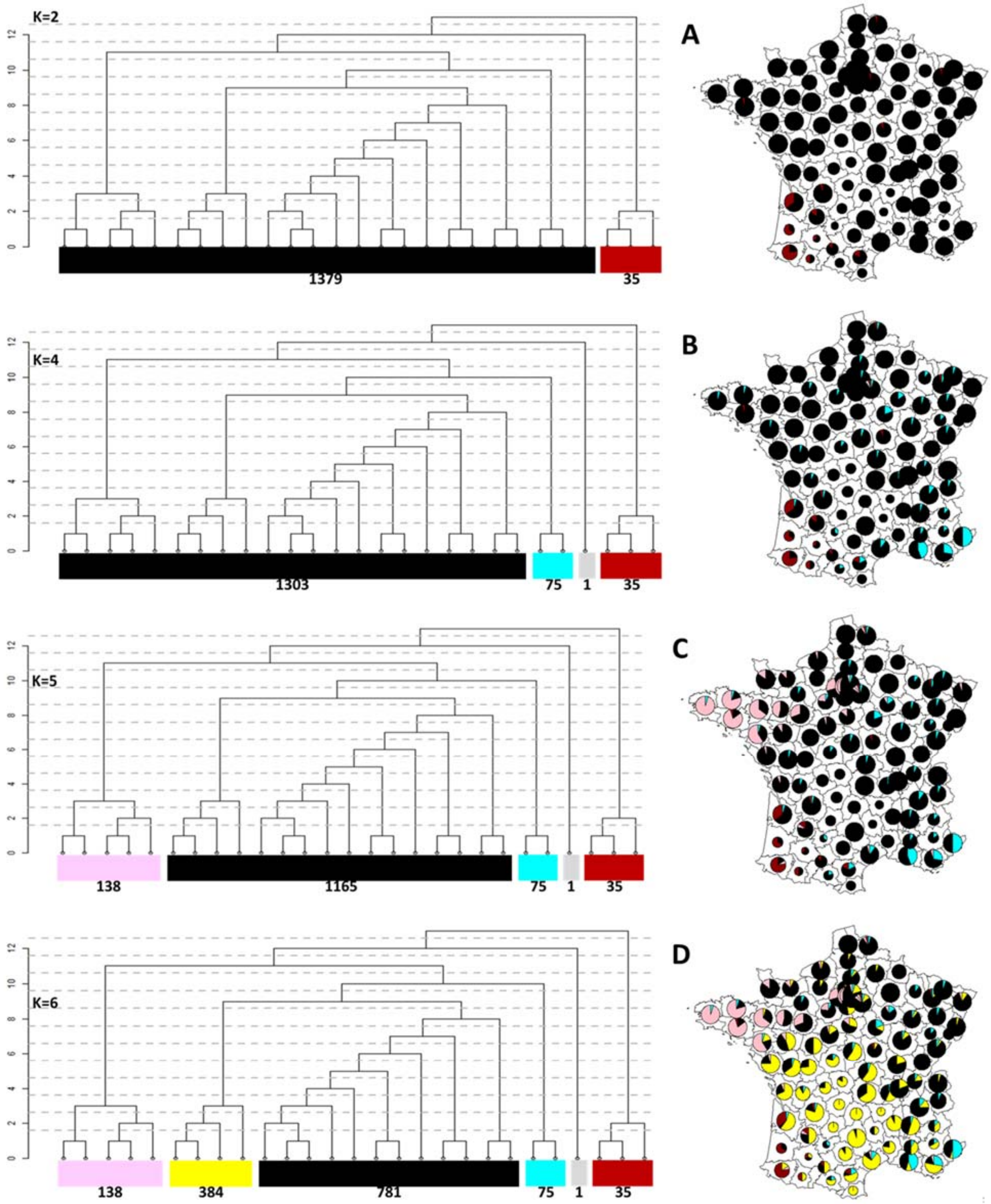


Figure S4: Genetic clusters in the SU.VI.MAX data inferred by the FineSTRUCTURE analysis at levels of the hierarchical clustering varying from k=2 (A), k=4 (B), k=5 (C) and k=6 (D).

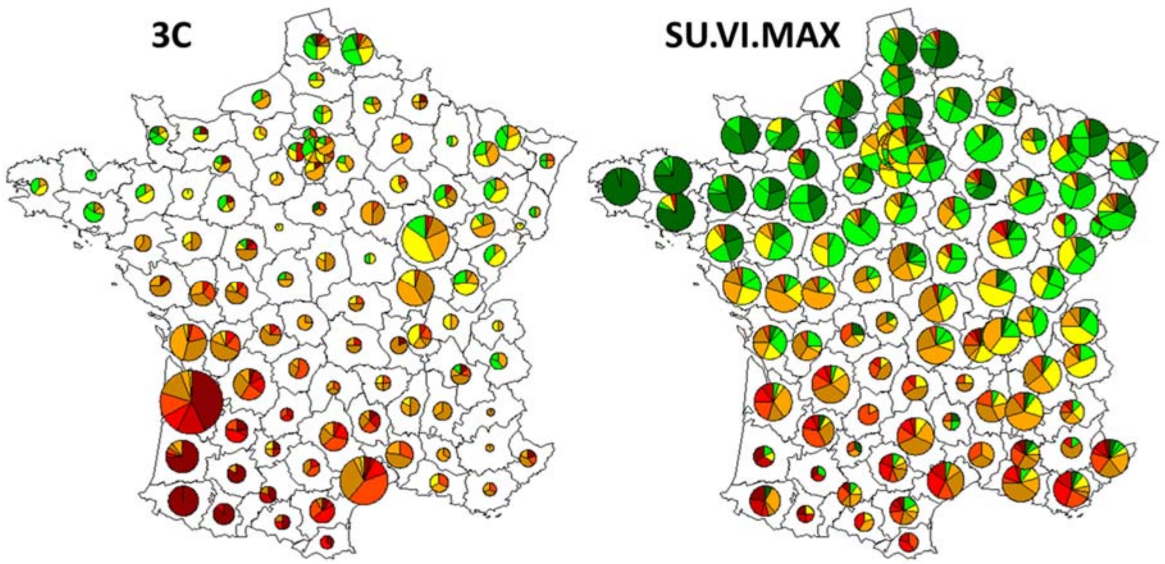


Figure S5: Distribution of PC1 values per *département* in the 3 Cities study and SUVIMAX. Colour of the points indicates the range of PC values. Red colours indicate negative values while green colours indicate positive values.

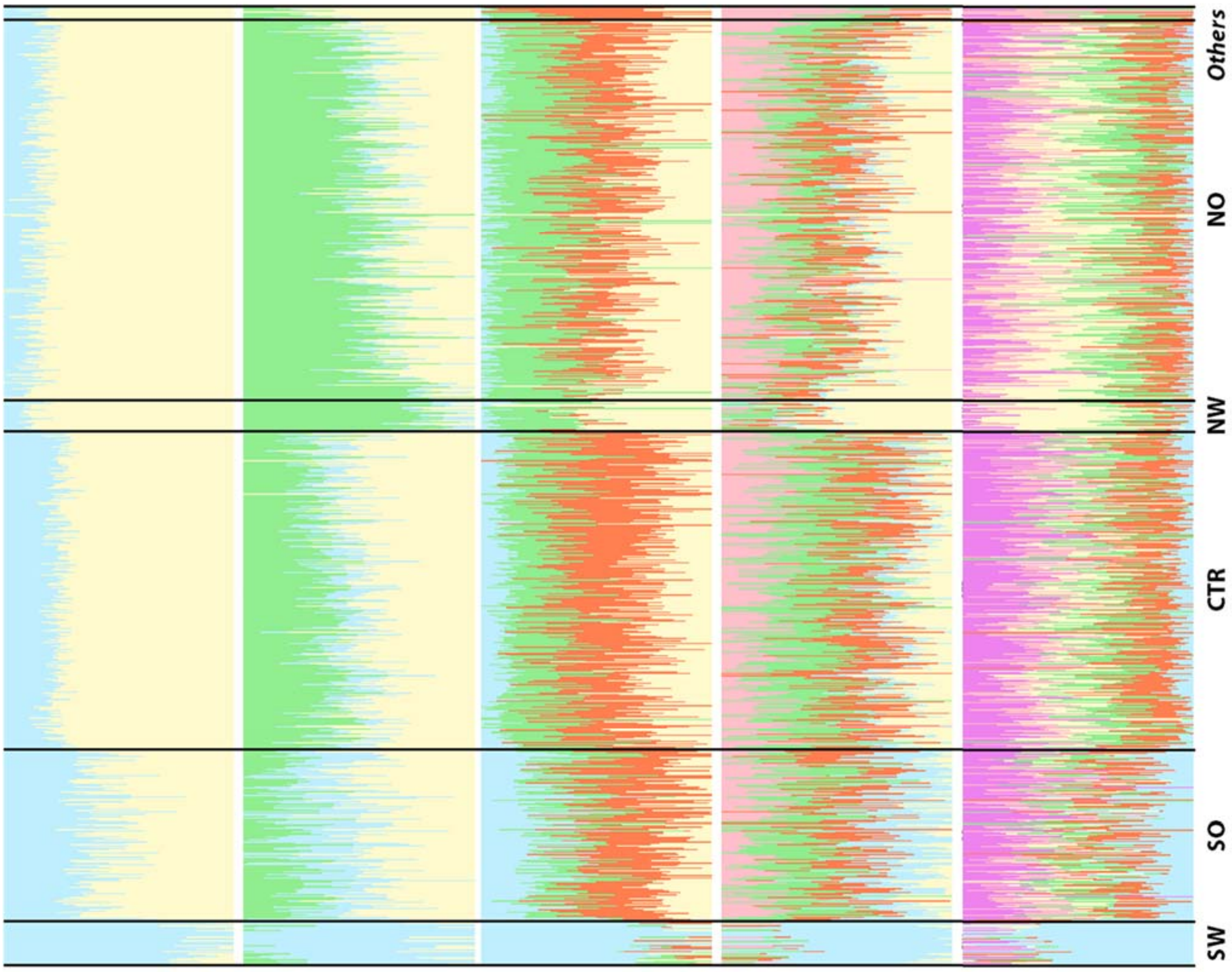


Figure S6: Results of ADMIXTURE assuming k=2 to k=6 in 3C. Each vertical line represents an individual and the different colors represent various ancestry components. In each cluster, individuals are sorted according to their geographical latitude.

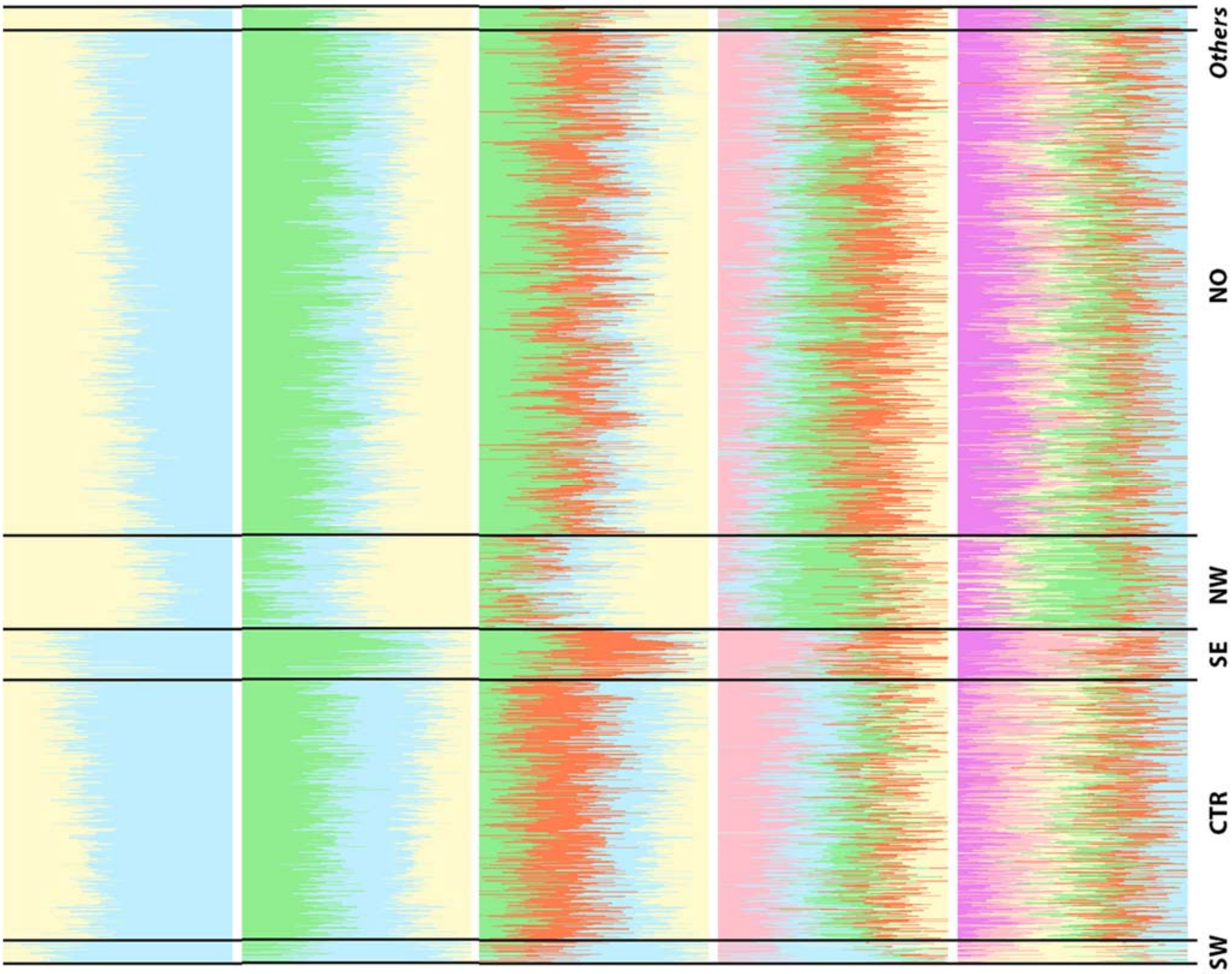


Figure S7: Results of ADMIXTURE assuming k=2 to k=6 in SU.VI.MAX. Each vertical line represents an individual and the different colors represent various ancestry components. In each cluster, individuals are sorted according to their geographical latitude.

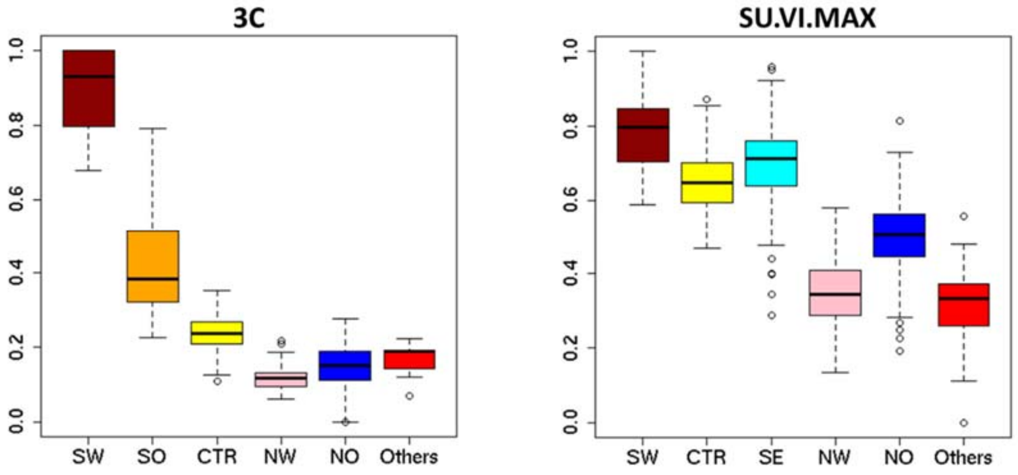

Figure S8: Proportion of the light blue ancestry at k=2 from ADMIXTURE in the 3 Cities study (left) and SU.VI.MAX (right)

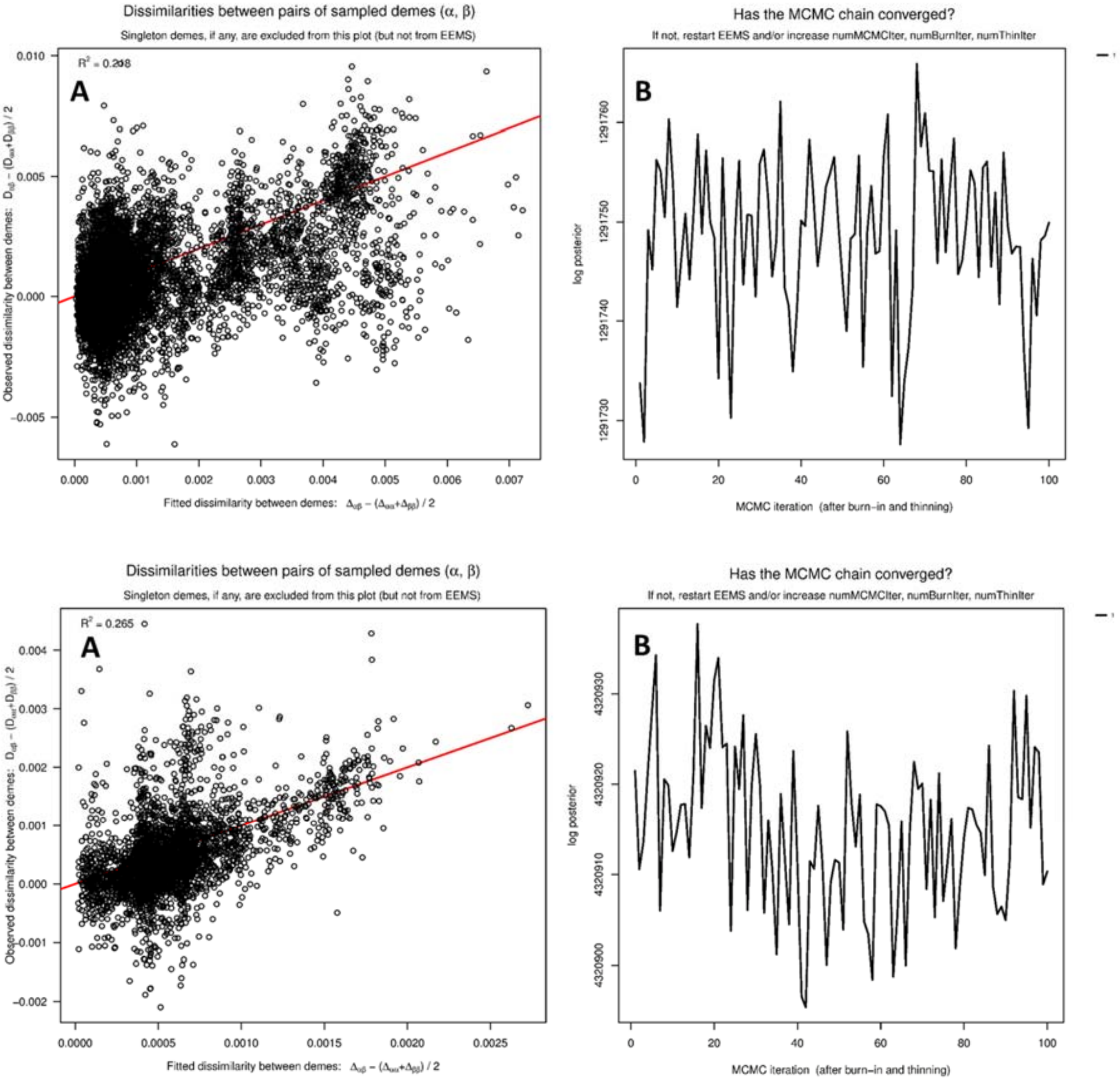


Figure S9: Estimated Effective Migration Surface Diagnostic Plots from EEMS performed on a grid of 250 demes with 770 3C (top) and 1,414 SU.VI.MAX individuals (bottom). (A) The observed versus expect dissimilarity between pairs of demes. Strong deviations from the fitted line (red) indicate pairs of demes much more genetically distant than expected. (B) The posterior probability log of the EEMS run, indicating whether the MCMC chains have converged.

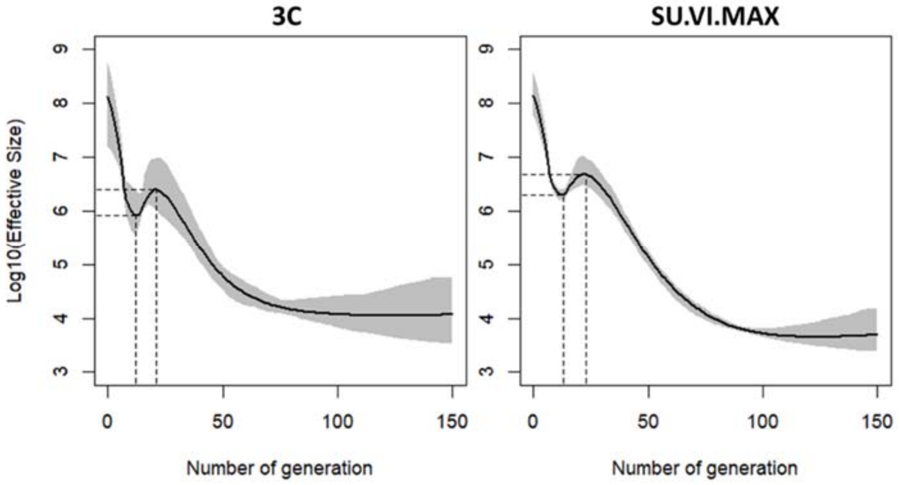


Fig S10: Evolution of effective size in 3C and in SU.VI.MAX.

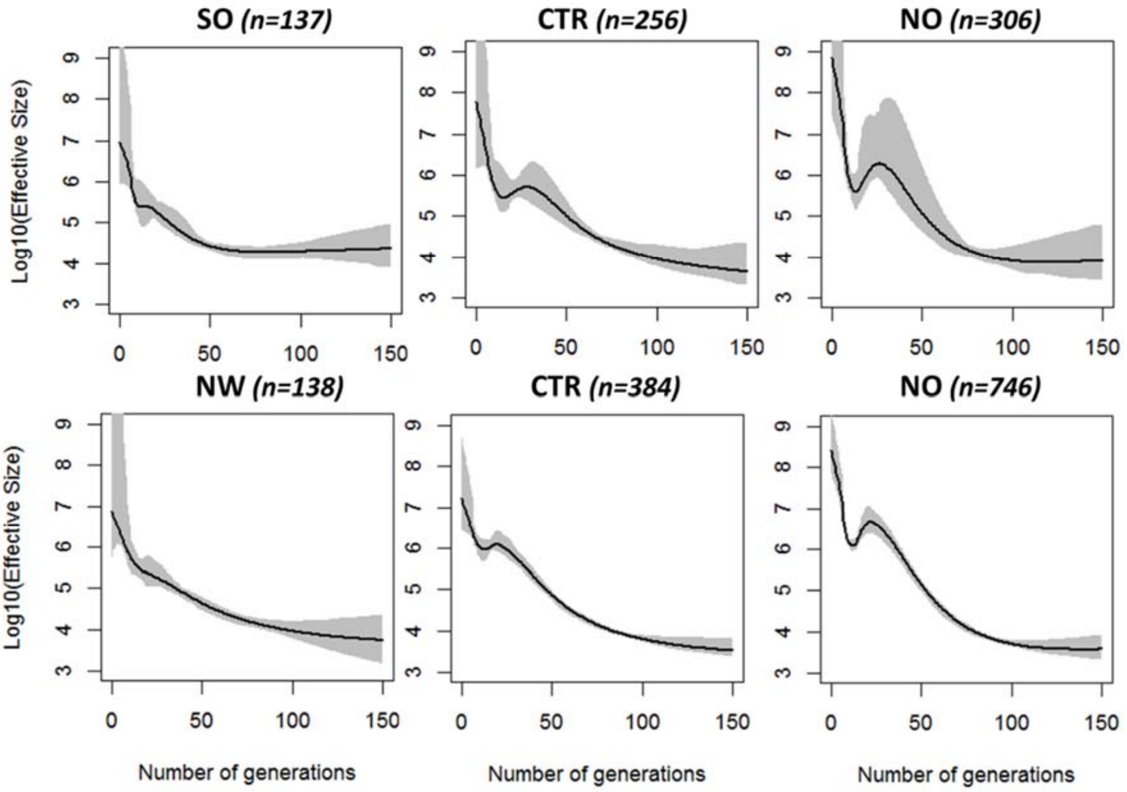


Figure S11: Evolution of effective size within cluster of France higher than 100 individuals in 3C (top: SO, CTR and NO) and in SU.VI.MAX (bottom: NW, CTR and NO).

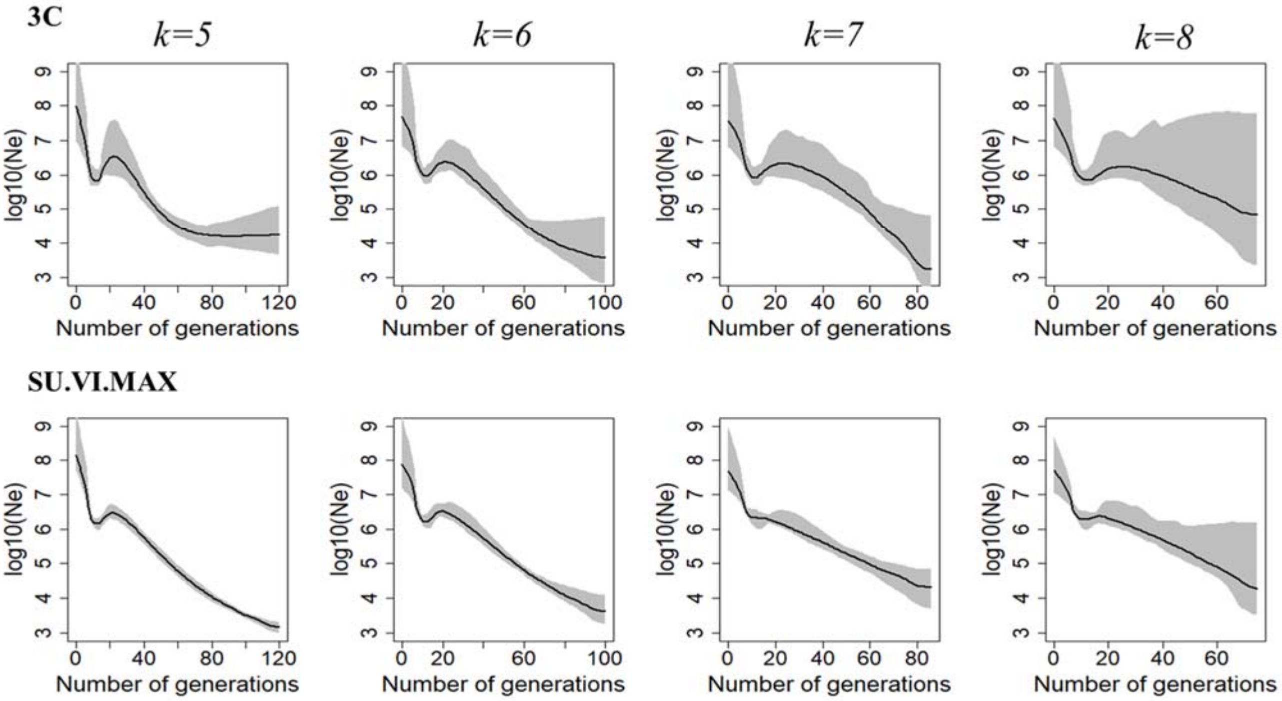


Figure S12: Evolution of effective size in the NO cluster with various mincm (k=5, 6, 7, 8) parameter in 3C (top) and in SU.VI.MAX (bottom).

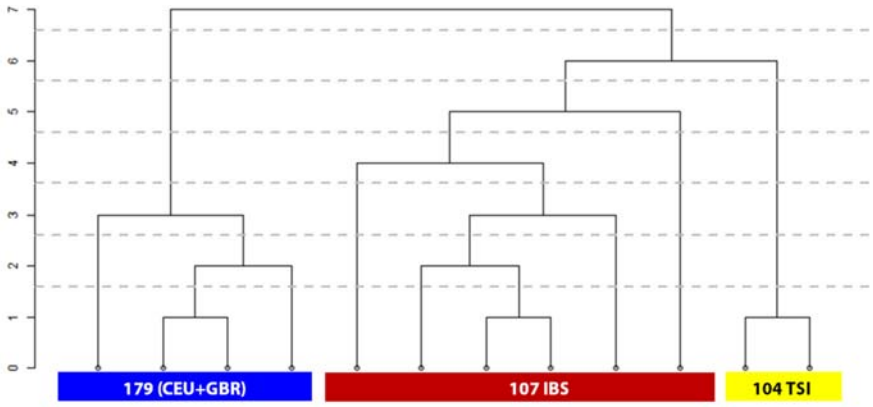


Figure S13: FineSTRUCTURE clustering of the European 1000G populations (390 individuals). The set of common SNPs with the 3C sample was used in this analysis.

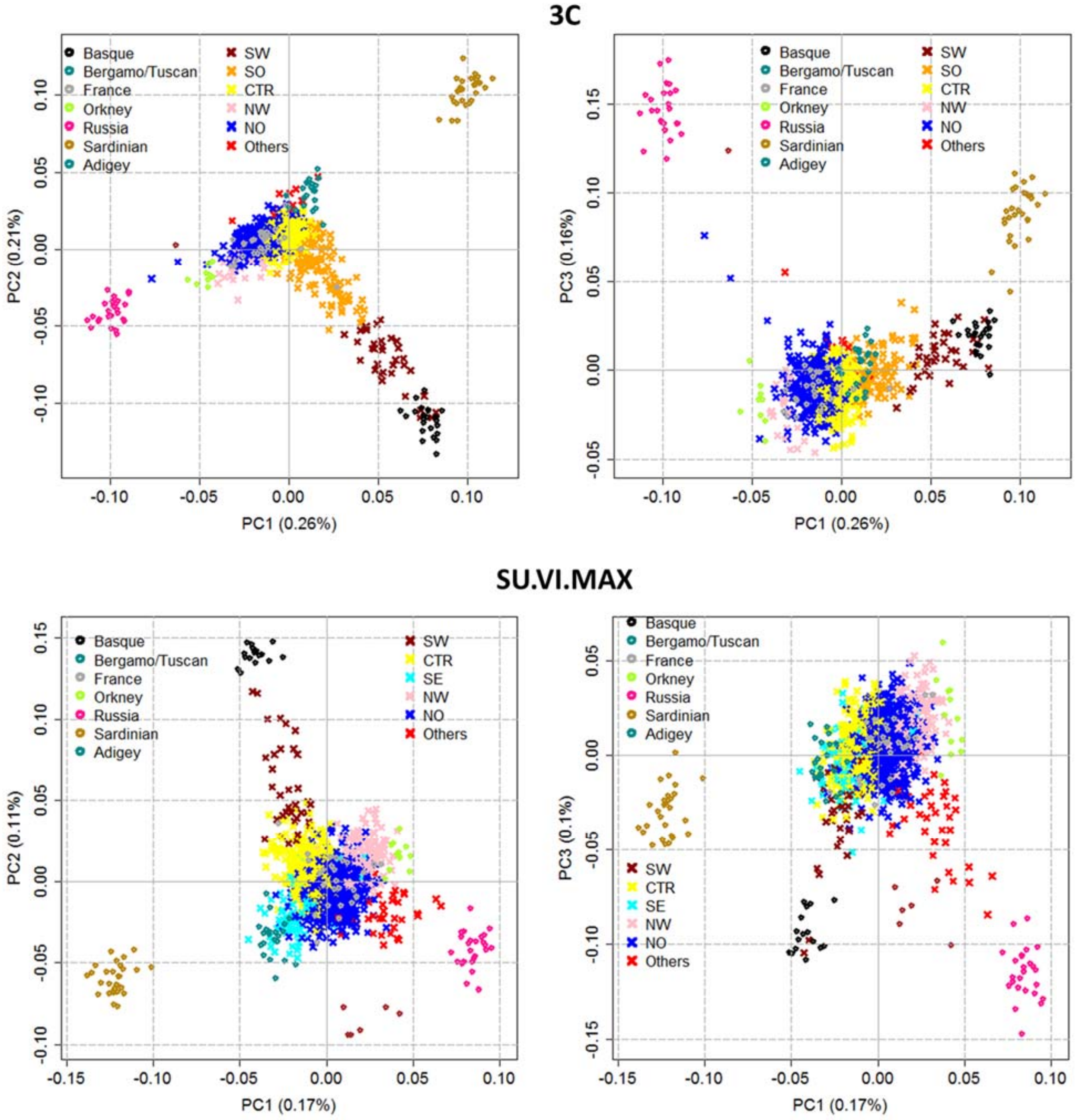


Fig S14: The scatter plot of the first three PCs from PCA performed on individuals from the 3 Cities study (top) or SU.VI.MAX (bottom) coloured according to the FineSTRUCTURE clustering and combined with the Europeans populations from the HGDP panel.

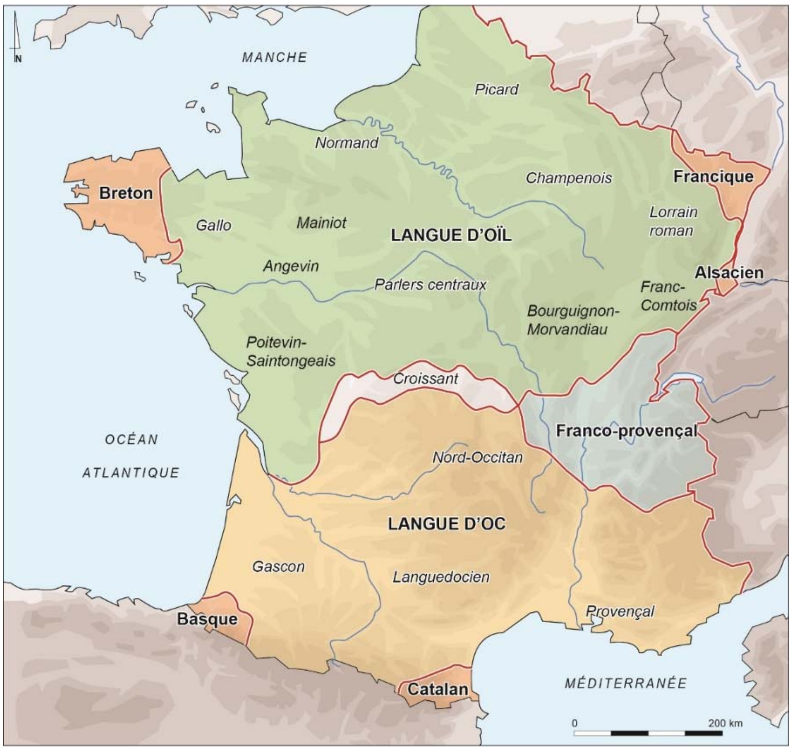


Figure S15: Map of roman, Germanic and other languages in France. The historical map has been plotted by © M. Monteil, Nantes University, UMR 6566 CReAAH.

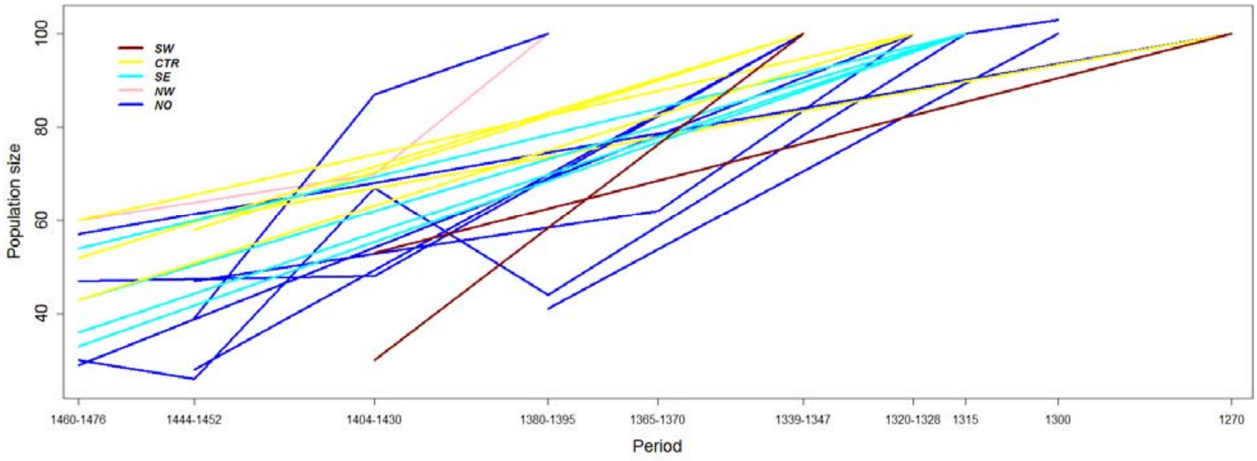


Figure S16: Change in population size in France between 1270 and 1476.

The table compiles results from more than 20 studies of local demography and is published in "*La France de la guerre de Cent Ans, Paris, Belin, 2009, rééd. 2013, p. 277-309* (chap. 8 - Les épidémies et la saignée démographique (XIVe-XVe siècles)".

Each trajectory represents the decay in population size of a French province. Lines are colored according to the geographical cluster where each province belongs to: NO: Artois, Cambrésis, Faucigny, Normandie orientale, Île-de-France, Champagne méridionale, Verdunois, Pays de Beaune. NW: Britanny. CTR: Languedoc, Limousin, Forez, Lyonnais, Dauphinois. SE: Basse Provence centrale, Basse Provence occidentale, Provence orientale, Haute Provence. SW: Bigorre, Navarre.
